# Metabolic integration of spectral and chemical cues mediating plant responses to competitors and herbivores

**DOI:** 10.1101/2022.09.02.506388

**Authors:** Alexander Chautá, André Kessler

## Abstract

Light quality and chemicals in a plant’s environment can provide crucial information about the presence and nature of antagonists, such as competitors and herbivores. Here we evaluate the roles of three sources of information - shifts in the red:far red (FR) ratio of light reflected off of potentially competing neighbors, induced metabolic changes to damage by insect herbivores, and induced changes to volatile organic compounds emitted from herbivore–damaged neighboring plants - to affect metabolic responses in the tall goldenrod, *Solidago altissima*. We address the hypothesis, that plants integrate the information available about competitors and herbivory to optimize metabolic responses to interacting stressors by exposing plants to the different types of environmental information in isolation and combination. We found strong interactions between the exposure to decreased Red:FR light ratios and damage on the induction of secondary metabolites (volatile and non-volatile) in the receiver plants. Similarly, the perception of VOCs emitted from neighboring plants was altered by the simultaneous exposure to spectral cues from neighbors. These results suggest that plants integrate spectral and chemical environmental cues to change the production and perception of volatile and non-volatile compounds and highlight the role of plant context-dependent metabolic responses in mediating population and community dynamics.

**Highlight:** Plants integrate spectral and chemical environmental cues to differentially induce production of volatile and non-volatile compounds

## Introduction

The ability to perceive, process, and integrate information from the environment is essential for any kind of behavioral or phenotypically plastic response and thus for the fitness of any organism (Chen *et al*., 2004). Plants are not an exception and, much like animals, have been shown to perceive and process information coded in light (Carvalho *et al*., 2011), sound (Khait *et al*., 2019), infochemicals (e.g. volatile organic compounds (VOCs) (Heil and Ton, 2008; Kessler, 2015), and touch (Mishra, Ratnesh Chandra Bae, 2019). Out of these, the perception of light and VOCs have received increased recent attention as they can encode information about the most important antagonistic interactions plants can have with other organisms, competition, and herbivory/pathogen attack, respectively (Karban, 2008). Plants perceive light with several specialized pigment molecules, such as chlorophylls and phytochromes. Among them, phytochromes regulate different processes such as germination, etiolation, shade avoidance, floral induction, induction of bud dormancy, tuberization, tropic orientations, and proximity perception (Smith, 1995; De Wit *et al*., 2013). Phytochromes are present in two interconvertible forms: P_r_ and P_fr_, and the relative cytosolic concentrations of P_r_ and P_fr_ are determined by environmental light quality (Rockwell *et al*., 2006). A low ratio of Red: Far-Red (R: FR) light transforms phytochrome into its inactive form (P_r_), which attenuates the degradation of PIF (phytochrome interacting factors), which, in turn, leads to different physiological changes in the plant. Most importantly for this study, the perception of elevated FR proportions, as reflected off of green leaves, allows plants to perceive potential competitors in their vicinity and preferentially allocate resources to competition by increasing elongating growth (Casal *et al*., 1987). Moreover, experiments on tobacco and tomato have demonstrated that this phytochrome-mediated perception of changes of the R: FR light ratio is also associated with a simultaneous, reduced allocation of resources into direct resistance to herbivores and an attenuation of induced chemical resistance (Izaguirre *et al*., 2006; Cortés *et al*., 2016). Likely underlying these attenuated metabolic responses to herbivory with corresponding effects on pathogen as well as herbivore resistance are apparent alterations of jasmonic acid (JA) and salicylic acid (SA)-mediated gene expression in plants exposed to elevated FR light ratios (De Wit *et al*., 2013; Fernández-Milmanda *et al*., 2020). Although, the direction of the effect of FR light on individual VOCs (Colquhoun *et al*., 2013; Kegge *et al*., 2015; Carvalho *et al*., 2016), and non-volatile compounds (Tegelberg *et al*., 2004; Kuo *et al*., 2015) and so the expression of different types of resistance (e.g. direct vs indirect resistance) may vary, the plants’ ability to perceive changes in the R: FR light ratio seems to allow a fine-tuning of the allocation of resources into competition or anti-herbivore defenses (Leone *et al*., 2014).

Like the perception of differences in light quality, the ability of plants to produce and perceive chemical environmental cues, such as VOCs seems to play an important role in coping with multiple environmental challenges. VOCs are crucial in mediating plant direct and indirect resistance against herbivores (Dudareva *et al*., 2013). After damage by herbivores plants emit increased and attacker-specific blooms of VOCs, often called herbivory-induce plant volatiles (HIPV)(Becker *et al*., 2015). These HIPVs are often repellent to foraging herbivores (direct repellence/ resistance) but can also function as effective cues that attract natural enemies of herbivores, such as predators and parasitoids (information-mediated indirect defenses) (Dicke and Baldwin, 2010; Becker *et al*., 2015). Moreover, HIPVs can also be perceived by other plants, which respond by priming or directly inducing increased production of defense-related secondary metabolites and so increased resistance in anticipation of oncoming herbivores (Karban *et al*., 2011; Okada *et al*., 2015; Morrell and Kessler, 2017; Kalske *et al*., 2019; Karban, 2021). The mechanisms of plant VOC perception are debated to this date (Erb, 2018) and may include direct alteration of membrane potentials (Heil, 2014), specific receptors (Gallie, 2015), and the transformation of VOCs into direct defensive compounds by the receiver plants (Sugimoto *et al*., 2014). However, very much like shifts in R: FR light ratios encode potential competition with neighbors, HIPVs provide a reliable cue for the probability of future herbivory. The fact, that plants have these different abilities to adaptively respond to changed light quality and HIPVs from neighbors, in turn, raises the question of how plants integrate these two different types of information to optimize responses to two of the most fitness-impacting environmental factors, competition, and herbivory.

*Solidago altissima* dominates early succession habitats in open environments in northeastern North America (Etterson *et al*., 2008; Howard *et al*., 2018). This species grows in dense patches where it competes for light with a diverse Astereacea-dominated plant community. Additionally, this species is attacked by a large diversity of insect herbivores (Maddox and Root, 1987, 1990). Most importantly, however, plant community composition (Carson and Root, 2000), as well as population genetic composition (Bode and Kessler, 2012; Uesugi and Kessler, 2013), are driven by a strong interaction between competition and insect herbivory on the dominant species S. altissima. Moreover, previous studies have demonstrated that *S. altissima* plants strongly respond to HIPVs from neighboring plants by priming and directly inducing changes in secondary metabolism and resistance (Morrell and Kessler, 2017). Moreover, HIPV-mediated plant-to-plant information transfer affects herbivore distribution (Rubin *et al*., 2015) and, in consequence is under strong herbivory-mediated natural selection (Kalske *et al*., 2019). The particularly strong interaction observed between competitive ability and herbivory in determining the persistence of *S. altissima* plants in a population and community, raises the more general question of how plants can utilize environmental information to adjust their phenotypes to varying environmental conditions while minimizing the combined, often synergistic impact of antagonistic biotic factors. The hypothesis that we are addressing here is that plants can integrate the information available on future herbivory and competition to induce metabolic changes that minimize the negative fitness effects of multiple interacting antagonists. Here we test two major predictions associated with this hypothesis: A) Secondary metabolite responses to herbivory should be altered in the presence of a neighbor (i.e. perception of lower R: FR ratio). B) Perception of oncoming herbivory (i.e. HIPVs from damaged neighbors) should be altered by the presence of a potentially competitive neighbor (i.e. perception of lower R: FR ratio). This hypothesis seems particularly relevant in the study system we chose for this project, the tall goldenrod *Solidago altissima*, where herbivory can be the major factor mediating competition with neighbors (Carson and Root, 2000; Uesugi and Kessler, 2013), and more nuanced and integrated responses to the combined perception of competitors and herbivores may maximize plant fitness. Here we use *S. altissima* in factorial manipulative experiments to address the above-mentioned hypothesis and further our understanding of how plants integrate two different sources of biotic environmental information (light and HIPVs).

## Materials and Methods

### Plant material

Seeds of *S.altissima* were bulk-collected in winter 2020 from plants around Bebe Lake, Ithaca, NY, and then stored in a freezer at −20°C. After a month, the seeds were put into LM1 germination mix soil (Lambert) for germination at Cornell University’s greenhouse with a photoperiod of 16:8 light: dark. Once the plants had germinated, they were repotted into individual clear polyethylene terephthalate plastic cups of 500mL capacity, and an initial measurement of the length of the plant and the number of leaves was taken. All plants were grown under high-pressure Sodium lamps that produce 200 μmol/m^2^/sec of white light in a photoperiod day-night 16:8. In addition, half of the plants were under supplemental Far-Red (FR) light, using an FR lamp (Forever Green Indoors, 730 nm) of 114 cm with led 32 bulbs. The FR lamps were covered with a blue filter (Roscolux, Supergel, Cinegel no. 83 Medium Blue) to remove residual red light following the protocol of Fernández-Milmanda *et al*. (2020). The lamp was located at 15 cm to the side of the plant and 10 cm from the ground to simulate the angle of light and the intensity coming from a neighboring plant. After one week, the second measurement of height and number of leaves was recorded to assess the effect of increased ratios of FR light on plant growth. After these measurements, two larvae of *Spodoptera frugiperda* in their third instar were added to each of 10 plants in each light treatment (FR and control), completing four groups of plants: Control, Damage, FR, and FR + Damage (Fig. 1A). After another four days with the larvae actively feeding. At this point, 10 plants in the damage treatment and 10 plants in the control treatment were used as emitter plants in a plant VOC-exposure experiment. Their VOC emissions were pulled into receiver plant chambers that included either control plants under normal light conditions or plants supplemented with FR light (Fig 1B). The chambers of both the emitter and receiver plants were connected through 0.7 cm diameter silicon tubing (BIO-RAD), and the chamber of the receiver was connected to an active air sampling vacuum pump (IONTIK) pulling air at about 450 ml/min. The pumps generate a constant flow of air from the emitter to the receiver plants (Fig. 1C). The pumps were changed twice a day, to ensure that there would be at least 22 hours of flow per day. After four days of VOC exposure, we collected VOCs using adsorbent traps and leaf material to analyze non-volatile metabolites. Volatile samples of each plant were taken by enclosing the plant into 500 mL polyethylene cups that were connected to an ORBO-32 charcoal adsorbent tube (Supelco®). The air was pulled through the charcoal traps using an active air sampling vacuum pump (IONTIK) pulling air at about 450 ml/min. Additionally, leaf samples were collected and flash-frozen in liquid nitrogen and later stored at −80°C until further analysis. To understand if the chemical response to damage is affected by the presence of a neighbor, we compared the volatile and non-volatile chemical profiles of the emitter plants (Control, Damage, FR, and FR+Damage). To understand if the perception of volatiles is affected by FR exposure, we compared the volatile and non-volatile, metabolites produced by the receiver plants.

**Figure 1.**
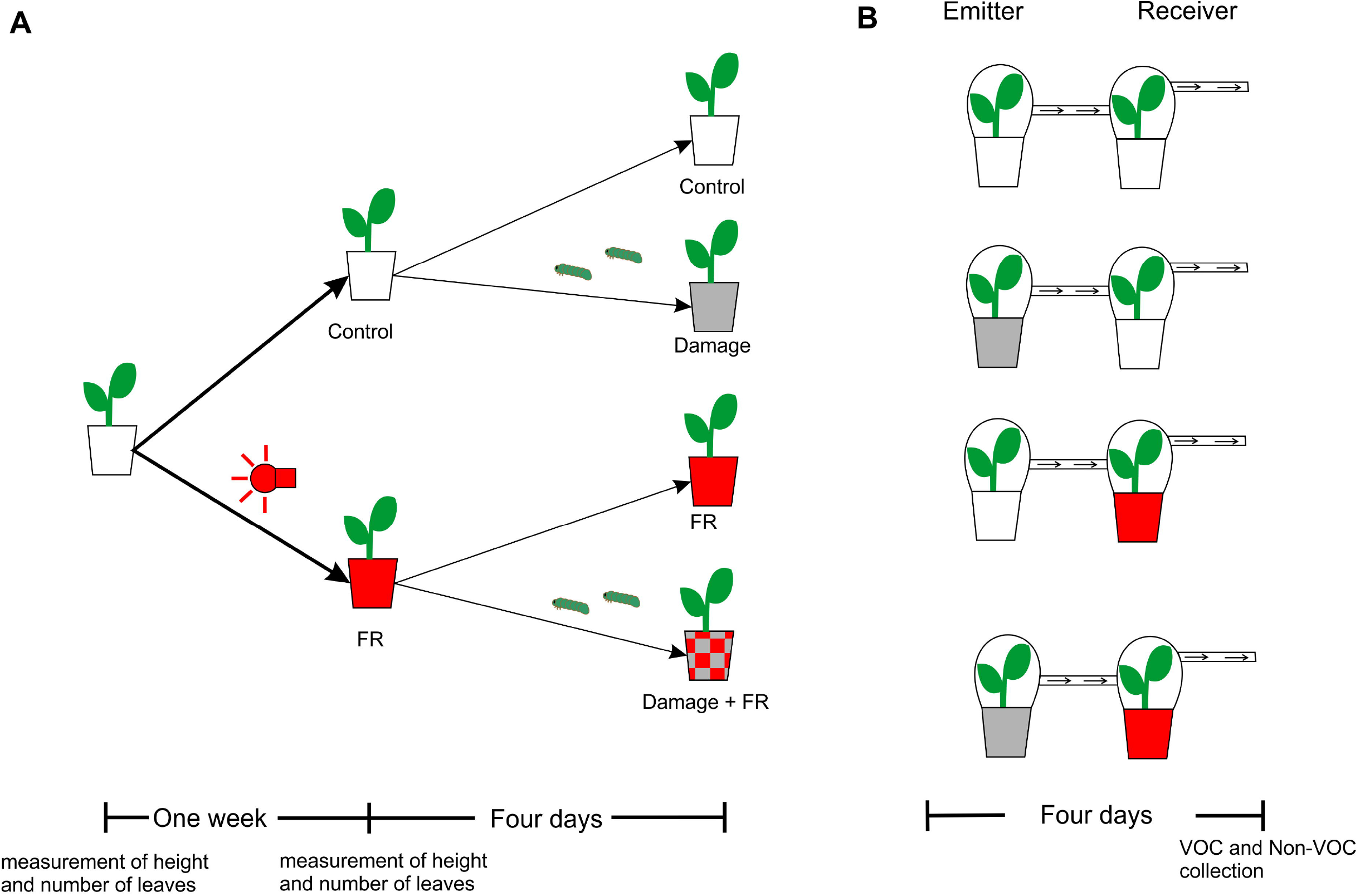
Experimental setup. **A)** Scheme showing the sequence of treatments of plants of *Solidago altissima*. Plants were divided into two groups; one was exposed to far-red (FR) light and the other was kept as a control. Then half of the plants in each treatment were damaged for four days by two *Spodoptera fruguiperda* larva in L2. **B)** Plants in the FR light and control treatments were set up to receive volatile organic compounds (VOC) from plants that were damaged by *S. fruguiperda*, for four days or from control plants that received no damage.

### Secondary metabolite analysis

For the analysis of VOCs emitted from the experimental plants, each of the ORBO-32 charcoal traps that were used in the collection was spiked with 5μL of tetraline (90 ng/mL) as an internal standard. The charcoal traps were washed with 400μL of dichloromethane, which was then injected in a Varian CP-3800 gas chromatograph (GC) coupled with a Saturn 2200 mass spectrometer (MS) and equipped with a CP-8400 autosampler. The GC-MS was fitted with a DB-WAX column, (Agilent, J&W Scientific) of 60 m × 0.25 mm id capillary column coated with polyethyleneglycol (0.25 mm film thickness). The temperature program began with an injection temperature of 225°C, heated from 45_C to 130°C at 10°C/minute, then from 130°C to 180 at 5°C/min, and finally from 180°to 250°C at 20°C/minute with a 5 min hold at 230 and 250°C. The samples were standardized by expressing signal intensity (peak area) of each peak relative to that of the internal standard. Compound and compound class identity were determined comparing mass spectra and retention time indices with NIST library records and previously published VOC data of *S. altissima* (Morrell and Kessler, 2017; Lawson *et al*., 2020).

For the high-performance liquid chromatography (HPLC) analysis of non-volatile compounds, leaf samples (150-250 mg/sample) were homogenized and extracted in 1 mL of 90% methanol using a FastPrep® tissue homogenizer (MP Biomedicals®) at 6 m/s for 90 s using 0.9-g grinding beads (Zirconia/Silica 2.3 mm, Biospec®). The samples were then centrifuged at 4 °C for 15 min at 14,000 rpm and analyzed 15 μL of the by HPLC on an Agilent® 1100 series HPLC. Using 99.9% acetonitrile and 0.25% H_3_PO_4_ as mobile phase. The elution system consisted of aqueous 0.25% H_3_PO_4_ and acetonitrile (ACN) which were pumped through a Gemini C18 reverse-phase column (3 μm, 150 × 4.6 mm, Phenomenex, Torrance, CA, USA) at a rate of 0.7 mL/min with increasing concentrations of ACN: 0–5 min, 0–20% ACN; 5–35 min, 20–95% ACN; and 35-45 min, 95% ACN. The area of each peak was standardized by the mass of the leaf tissue extracted. The individual compound or class identity was determined based on the UV spectra and retention times of authentic standards.

### Statistical analysis

The differences in growth between FR and control plants were analyzed using a Students’ t-test. The overall composition of volatile and non-volatile plant secondary metabolites was inspected using nonmetric multidimensional scaling (Bray–Curtis distance matrix; metaMDS in *vegan* package) and tested for the effects of the FR light exposure and herbivory on the composition with a PERMANOVA with 999 permutations using the adonis2 function in the vegan package using the trials as strata. For the emitters, we used the exposure to FR and damage as independent factors, and the relative abundance of compounds as dependent factors. For the receivers, we used the exposure to FR and the damage on the emitter plant as independent factors, and the relative abundance of compounds as dependent factors. If PERMANOVAs had shown significant results, a *post hoc* test was run using the function *pairwise.adonis2* from the library *pairwise.adonis* (Martinez Arbizu, 2020). We adjusted the P values for the multiple corrections using the false discovery rate (FDR) adjustment (*p. adjust* in package *stats*). Additionally, individual ANOVAs were run for each volatile and non-volatile compound from emitter and receiver plants. Those that showed variation with treatments were included in a heatmap analysis for easier visualization of the complex differences. All statistical analyses were performed using the R program (R Team Core, 2021).

## Results

### Effect of RFR light on plant growth

Exposure to supplemental FR light resulted in increased stem elongation relative to plants under normal light conditions (*t* = −10.264, df = 63, p < 0.001, Fig 2A), however there was no effect on the number of new leaves that grew between measurements (*t* = −0.93743, df = 63, *p* = 0.3521, Fig. 2B).

**Figure 2.**
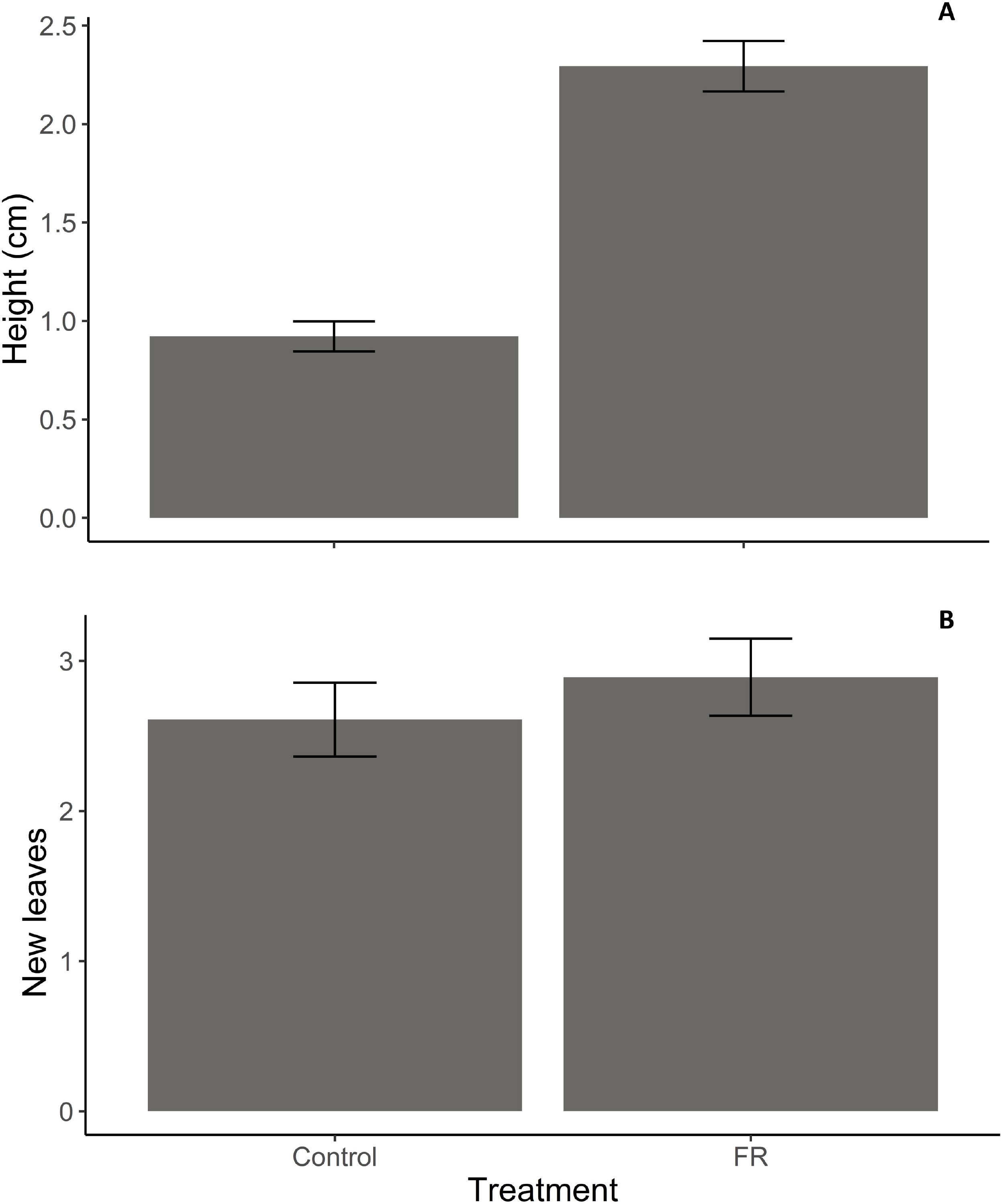
Plant growth responses to far-red (FR) light exposure. **A)** Mean (±SEM) stem height and **B)** Mean (±SEM) number of new leaves produced by *Solidago altissima* plants growing under regular light conditions supplemented with FR light (reduced red: far-red ratio, RFR) or under regular (control) light conditions over one week.

### Effect of herbivory and FR radiation on plant chemistry

Both FR exposure and herbivore damage influenced VOC bouquets (PERMANOVA, *F*_1,36_=9.5016, *p*=0.001 and *F*_1,36_= 4.3728, *p*=0.007 respectively) and we identified a strong interaction between both factors (FR x herbivore damage, *F*_1,36_= 7.2251, *p*=0.001). The Post hoc analyses identified differences in VOC bouquet compositions between all the treatments except between tplants damaged by herbivores and plants that were exposed to both, enhanced FR light and herbivore damage (FR+Damage) (Table 1). This result is also apparent in an NMDS analysis (Fig. 3A, stress value= 0.1150099). *Post-hoc* ANOVAs revealed 30 individual VOCs whose emissions varied with treatment and predominantly increased in response to herbivore damage or the combination of FR exposure and damage (Fig. 5A). Nonvolatile compound compositions were also affected by supplemented FR exposure (PERMANOVA, *F*_1,36_=3.2915, *p*= 0.011) but only marginally by damage (*F*_1,36_=2.0827, *p*=0.060), however, we observed a strong interaction between both factors (FR x herbivore damage, *F*_1,36_=5.4194, *p*=0.001).

**Table 1.** VOC composition as a function of light quality and herbivore damage. Pairwise (*post hoc*) comparison of the VOC bouquets emitted from *Solidago altissima* plants that had been exposed to control light (Control), or regular light supplemented with FR light (FR), while being not damaged or damaged by two larvae of *S. frugiperda* in L2 instar (Damage). *Represent statistical differences (p<0.05).

| Treatment | Control | Damage | FR |
| --- | --- | --- | --- |
| Damage | $F_{1,18} = 3.872, p = 0.001 *$ | | |
| FR | $F_{1,18} = 19.116, p = 0.001 *$ | $F_{1,18} = 7.847, p = 0.001 *$ | |
| FR+Damage | $F_{1,18} = 5.7041, p = 0.001 *$ | $F_{1,18} = 0.861, p = 0.283$ | $F_{1,18} = 7.7928, p = 0.001 *$ |

**Figure 3.**
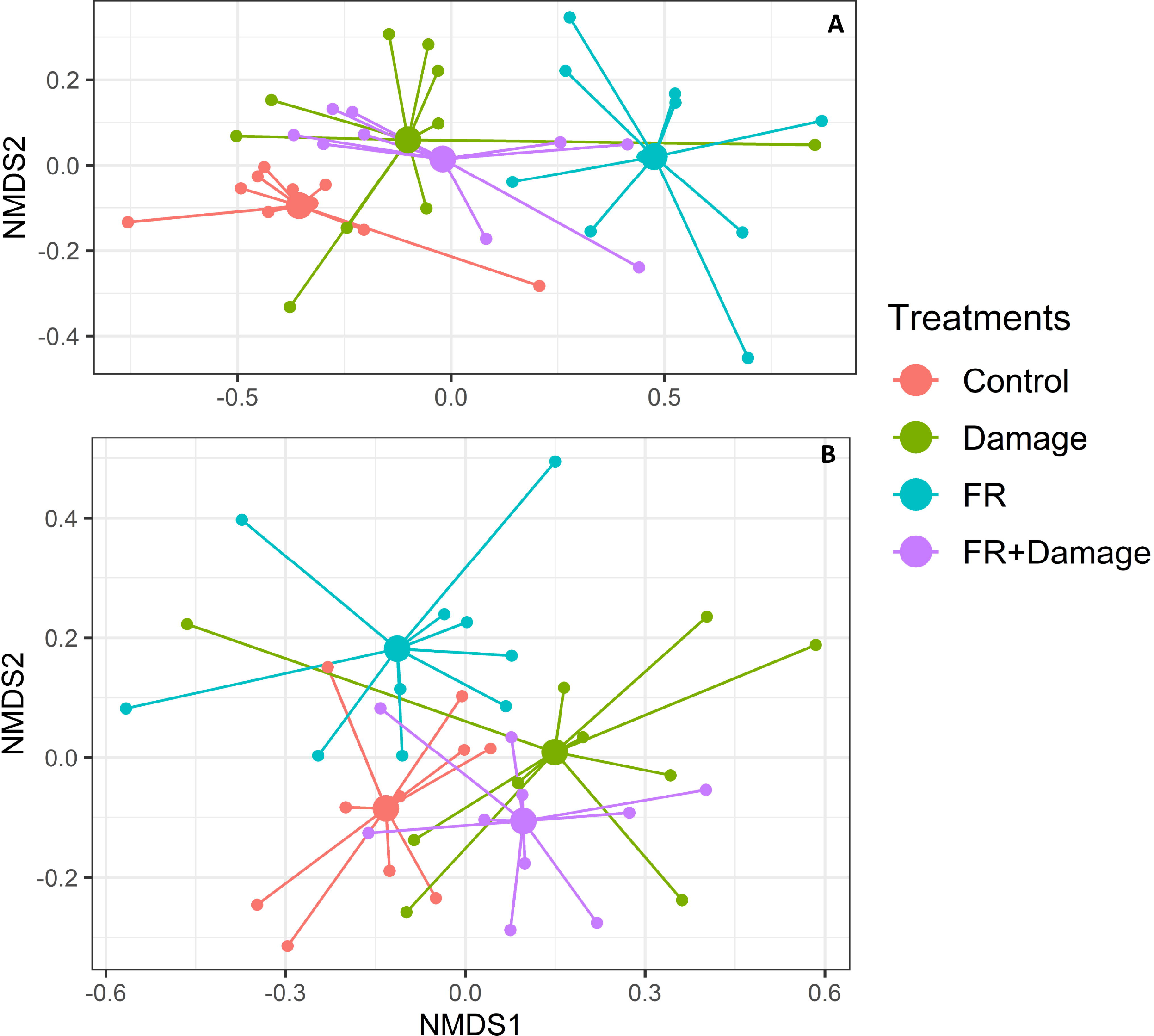
Plant secondary metabolite production in response to supplemented far-red (FR) light exposure and herbivore damage. Non-metric multidimensional scaling (NMDS) of **A)** volatile organic compound emissions (stress value=0.115) and **B)** non-volatile secondary metabolite production (stress value=0.163) of *Solidago altissima* plants, growing under reduced red: farred light ratios (FR), with damage by larvae of *Spodoptera frugiperda* (Damage), the combination of FR and damage (FR +Damage) or under control light (control) conditions.

**Figure 4.**
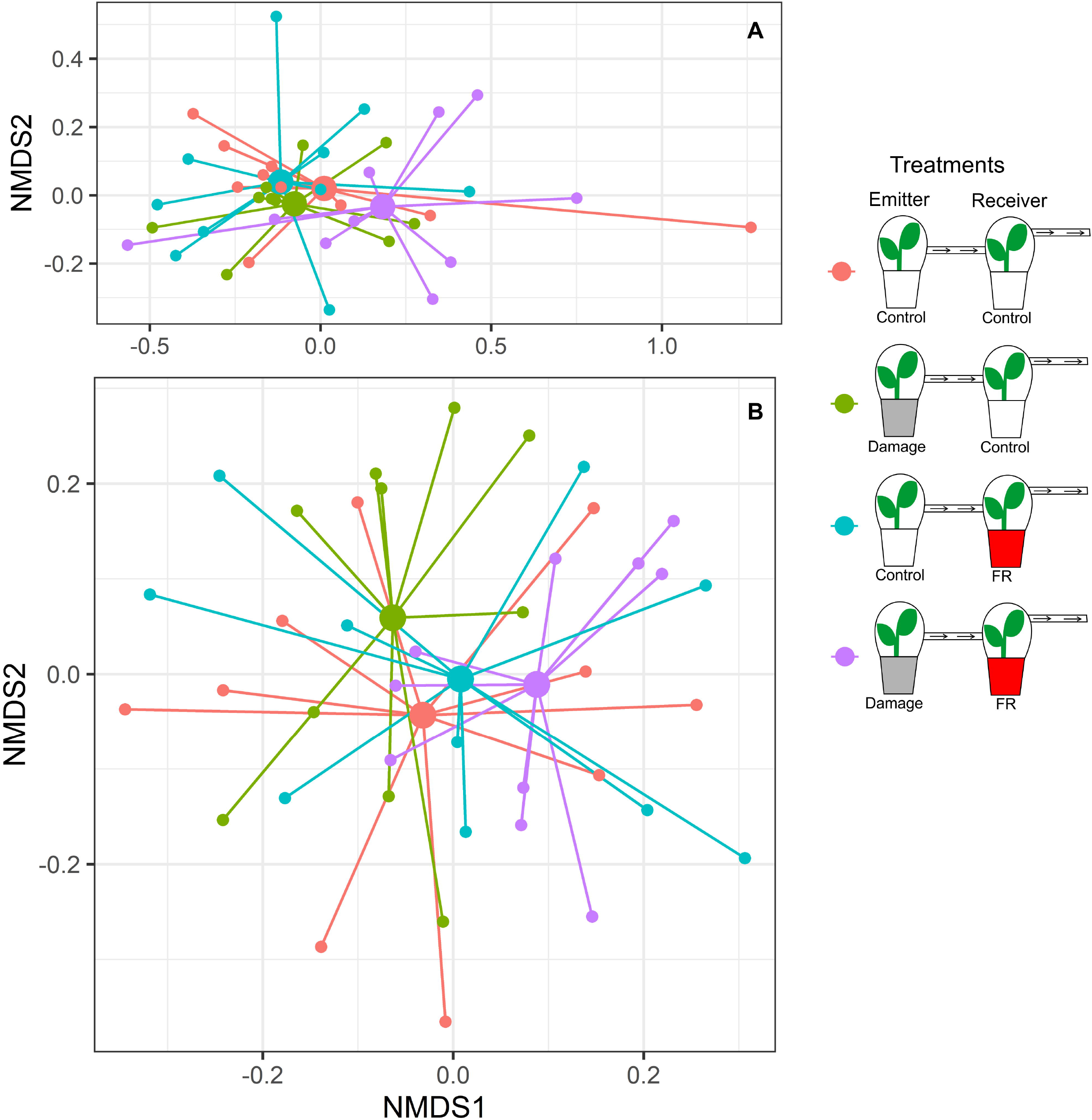
Plant secondary metabolite production in response to supplemented far-red (FR) light exposure and volatile organic compounds (VOCs) from neighbors. Plant secondary metabolite production of plants under supplemented far-red light (Red pots) or normal light (white pots), that were exposed to VOCs from control plants or plants that have been damaged by two larvae of S. furgiperda (grey pots). Non-metric multidimensional scaling (NMDS) of **A)** VOC emissions (stress value=0.08728402) and **B)** non-volatile secondary metabolite production (stress value=0.2796113) of *Solidago altissima* plants under the different exposure treatments.

**Figure 5.**
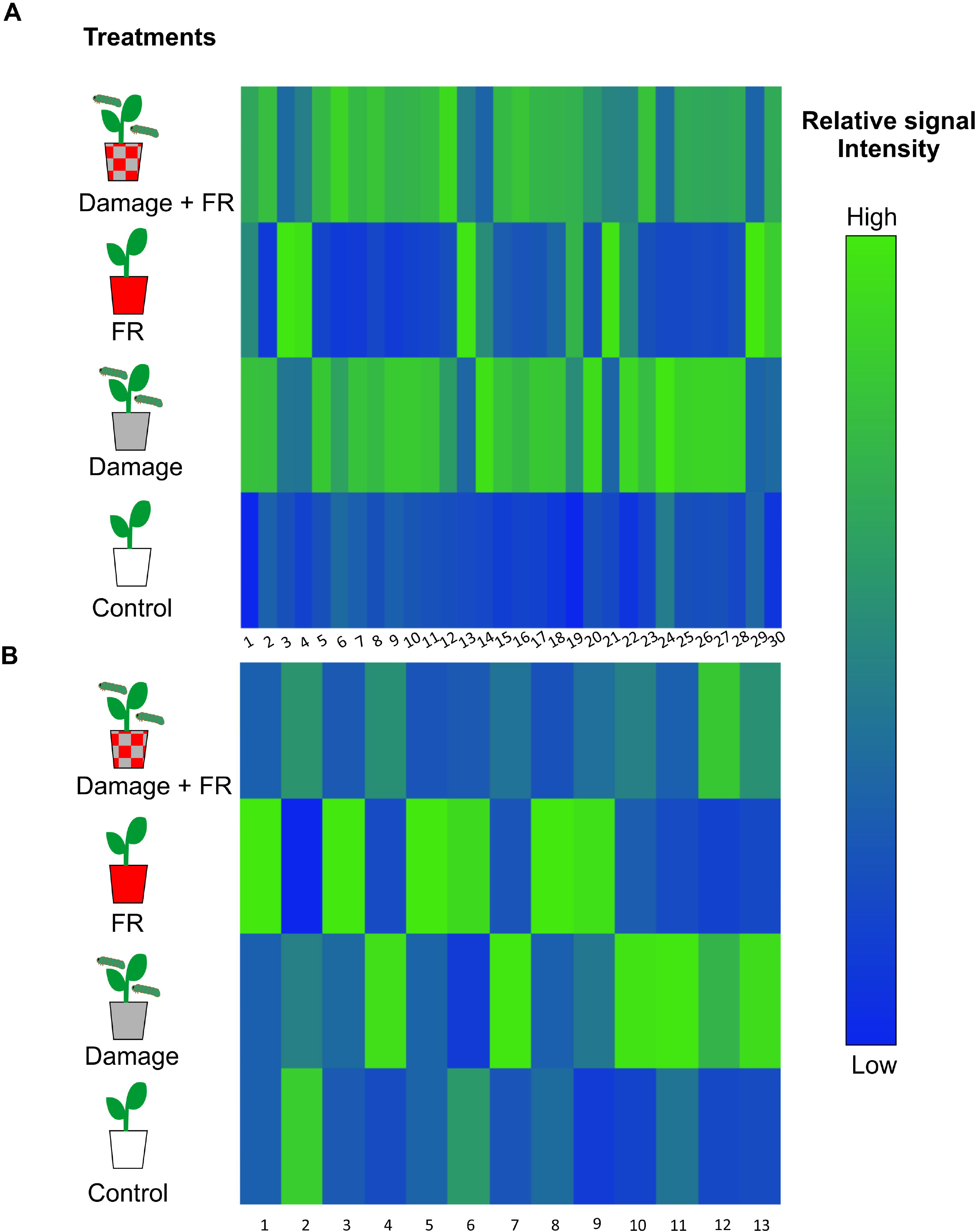
Figure 5. Differential induction patterns of individual secondary metabolites in response to supplemental far-red (FR) light exposure and herbivory. Heat map of (A) the emission of volatile organic compounds (VOCs) with tentative identification (**1.** Cymene, **2.** Dimethyl-1,3,7-nonatriene, **3.** (3Z)-Hexenyl acetate, **4.** 1,3,8 -Menthatriene, **5.** Unknown 1, **6.** Unknown 2, **7.** Unknown 3, **8.** p-Copaene, **9.** β-Bourbonene, **10.** α-Cubebene, **11.** Linalyl isobutyrate, **12.** Unknown 4, **13.** Bornyl acetate, **14.** 1,5-Cyclodecadiene, **15.** β-Caryophyllene, **16.** Unknown 5, **17.** Unknown Sesquiterpene 1, **18.** α-Humulene, **19.** Unknown Sesquiterpene 2, **20.** γ-Muurolene **21.** Unknown sesquiterpene 3, **22.** Methyl salicylate, **23.** Unknown 6, **24.** Unknown aliphatic compound, **25.** Unknown 7, **26.** Unknown 8, **27.** Unknown 9,. **28.** α-Phellandrene, **29.** β-pinene, **30.** Limonene) and (B) the production of non-volatile compounds (**1.** Unknonwn 10, **2.** Diterpene 1, **3.** Coumaric acid 1, **4.** Coumaric acid 2, **5.** Chlorogenic acid, **6.** Coumaric acid 3, **7.** Flavonoid 1, **8.** Diterpene 2, **9.** Diterpene 3, **10.** Diterpene 4, **11.** Diterpene 5, **12.** Diterpene 6, **13.** Diterpene 7) whose production is significantly varying with treatment (*p*<0.05). The different treatments include untreated controls, plants exposed to increased FR radiation (RFR), plants damaged by *Spodoptera frugiperda* caterpillars (Damage), and plants that received both treatments (FR + Damage). Different shades of color represent different intensity signal.

The *Post hoc* analyses of non-volatile secondary metabolite composition revealed differences between all the treatments except for the comparison between damage vs FR+Damage and Control vs FR+Damage (Table 2), which is also reflected in the NMDS analysis (Fig. 3B, stress value= 0.2587829). ANOVAs identified 13 non-volatile compounds that show a pronounced increase in two treatments (Damage and FR) but their productions tended to be lower in the combined FR+Damage treatment (Fig. 5B). From those compounds, seven are diterpene derivatives, two coumaric acid derivatives, one flavonoid, one chlorogenic acid derivative, and one currently unknown compound.

**Table 2.** Non-volatile secondary metabolite composition as a function of light quality and herbivore damage. Pairwise (*post hoc*) comparison of the non-volatile secondary metabolite composition *Solidago altissima* plants that had been exposed to control light (Control), or regular light supplemented with FR light (FR), while being undamaged or damaged by two larvae of *S. frugiperda* in L2 instar (Damage).

| Treatment | Control | Damage | FR |
| --- | --- | --- | --- |
| Damage | $F_{1,18} = 2.5411, p = 0.03 *$ | | |
| FR | $F_{1,18} = 7.8519, p = 0.003 *$ | $F_{1,18} = 3.7581, p = 0.001 *$ | |
| FR+Damage | $F_{1,18} = 1.33, p = 0.153$ | $F_{1,18} = 0.887, p = 0.461$ | $F_{1,18} = 4.7507, p = 0.002 *$ |

### Effect of FR on the perception of VOCs

The overall composition of VOCs emitted from plants exposed to VOCs from neighboring plants did not change with exposure to increased FR light ratios (*F*_1,36_= 1.444, *p*=0.095) or with the exposure to VOCs from damaged plants (*F*_1,36_= 1.459, *p*=0.106), neither was there an interaction between both factors (*F*_1,36_= 1.543, *p*=0.086. Fig. 4A). However, ANOVAs on individual compounds identified differential response patterns for 5 compounds. Four of those compounds were stronger emitted from plants that were exposed to increased FR light and received VOCs from control plants. However, those compounds showed lower emission rates when simultaneously exposed to increased FR light and VOCs from damaged neighbors, indicating an integration of the two types of environmental information (Fig. 6A). The fifth compound was emitted in higher amounts from plants exposed to FR light while also receiving VOCs from damaged neighbors (Fig. 6A).

**Figure 6.**
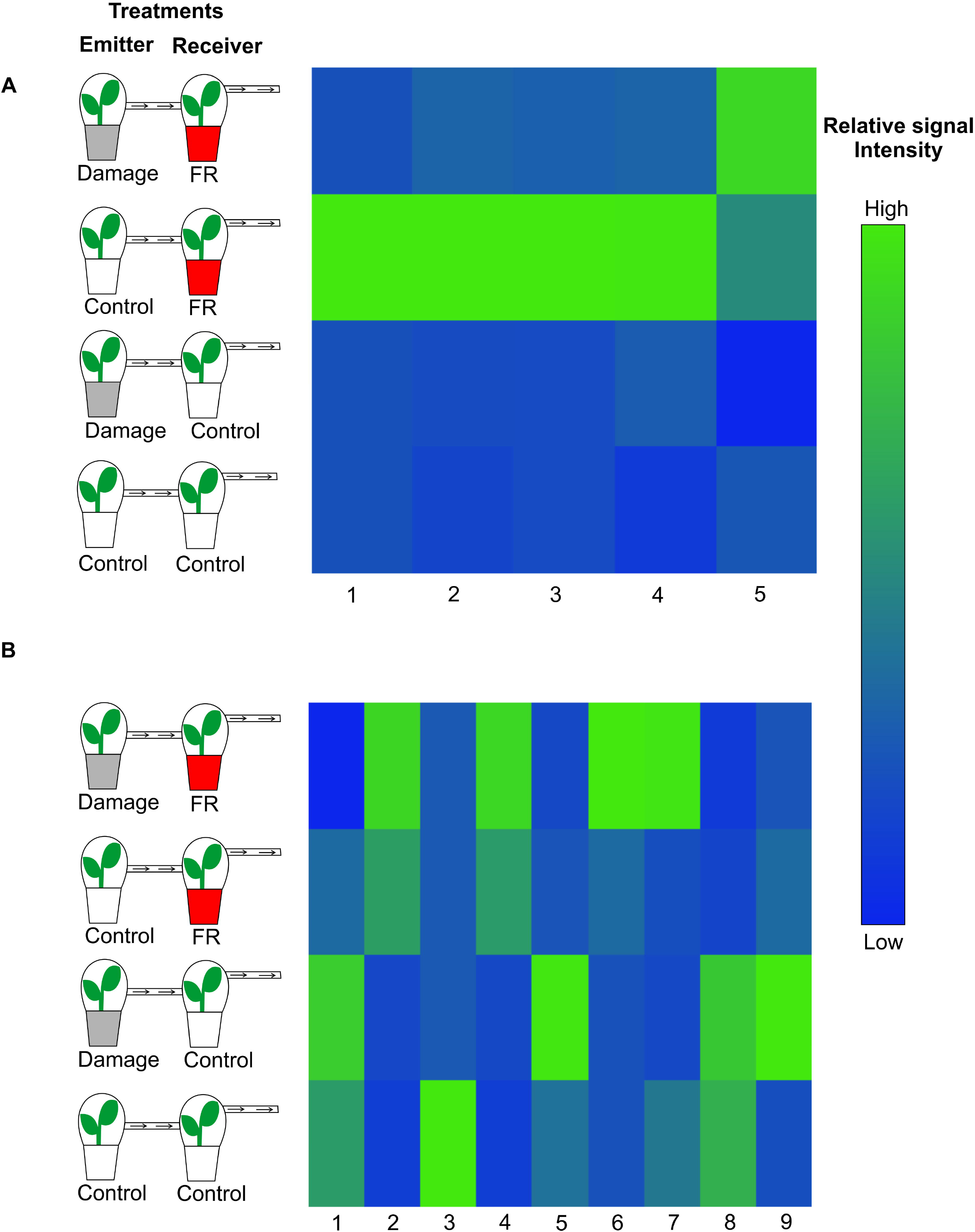
Differential induction patterns of individual secondary metabolites in response to spectral (far-red (FR supplementation) and volatile organic compound (VOC)-mediated information from neighboring plants. Heat map of **A)** the emission of VOCs with tentative identification (**1.** Unknown 1, **2.** α-Ylangene, **3.** Bornyl acetate, **4.** Unknown 2, **5.** β-Phellandrene) and **B)** the production of non-volatile compounds (**1.** Coumaric acid, **2.** Caffeic acid, **3.** Flavonol 1, **4.** Diterpene 1, **5.** Flavonol 2, **6.** Unknown, **7.** Chlorogenic acid, **8.** Diterpene 2, **9.** Diterpene 3) whose production significantly vary with treatment (*p*<0.05). The different treatments include plants exposed to supplemented FR radiation or regular light and plants exposed to VOCs from undamaged plants or VOC from damaged by *Spodoptera frugiperda* caterpillars, and plants that were exposed to both FR light supplementation and VOCs from damaged plants. Different shades of color represent different intensity signal.

Like the VOC responses, overall non-volatile compound composition from plants that were exposed to increased FR radiation or were exposed to VOCs from herbivore-damaged plants did not change (*F*_1,36_= 1.50148, *p*=0.119, *F*_1,36_= 0.85814, *p*=0.497 respectively) relative to that from controls. Moreover, perception of neighbors perception and herbivory on neighboring plants (FR and Damage VOCs) did not interact to affect non-volatile secondary metabolite production (*F*_1,36_= 0.96695, *p*=0.380, Fig. 4B). However, ANOVA analyses of individual compounds showed significant changes for some to them (Fig. 6B).

## Discussion

Plants can perceive neighbors by the shift in the R: FR ratio of light reflected off of green leaves and are commonly observed to respond with accelerated stem elongation (Demotes-Mainard *et al*., 2016). In confirmation of these earlier findings, we found *S. altissima* plants responding in a very similar way. The exposure of the plants to increased FR light radiation resulted in stem elongation but did not alter the number of leaves produced. At the same time, plant secondary metabolism as well as plant metabolic responses to herbivory and to VOCs from neighboring plants were strongly affected by FR perception. Specifically, *S. altissima* plants exposed to both, supplemented FR light, and herbivory induced differences in secondary metabolite production (volatiles and non-volatiles) with both factors interacting, suggesting that both, constitutive and induced secondary metabolite production are strongly affected by the exposure to spectral cues from neighboring plants. This coordinated change in growth and metabolism in response to perceived potential competitors has recently been interpreted as a differential allocation of resources into plant competitive ability, and defensive functions (Izaguirre *et al*., 2006; Leone *et al*., 2014) (Izaguirre *et al*., 2006; Leone *et al*., 2014). Similarly, but to a somewhat smaller extent, the ability of plants to perceive volatiles coming from neighboring plants (with and without damage) was affected by the exposure to FR light. These findings are significant as they suggest that plants process information about actual herbivory as well as potential future herbivory differently when exposed to neighboring plants and plants differentially integrate information about different types of antagonists (e.g. herbivores and competitors) to induce metabolic responses.

### FR light and plant growth

In shade-intolerant plants a low ratio of R: FR light induces differential growth such as stem elongation (Fankhauser and Batschauer, 2016) that can provide a competitive advantage over neighbors in the natural habitats (Demotes-Mainard *et al*., 2016). This stem elongation is regulated by gibberellin A1 and Indole-3-acetic acid (IAA)-mediated cell expansion rather than cell propagation (Kurepin *et al*., 2007; Pierik *et al*., 2014) and thus results in an increase in the internode distance rather than an increase in the number of internodes (Demotes-Mainard *et al*., 2016), *Solidago altissima* plants respond to FR supplementation in a similar way despite the fact that this species seems well adapted to growth in dense, high-competition environments. Similar plant-endogenous signaling mechanisms are thought to also mediate the correlated changes in plant secondary metabolism and the changes in the inducibility of metabolic responses to other environmental cues and stressors, such as herbivory (Izaguirre *et al*., 2006). Thus, the functional question for why plants induce changes to competition-indicating light quality has to be answered on two levels. On one hand, we need to explain why secondary metabolism changes in response to different light quality in the first place (i.e. the potential benefit of altered constitutive defenses). And, on the other hand, one has to probe the potential effects of light-quality-mediated changes on the perception of other stressors, such as herbivory (i.e. integration of environmental information from different sources).

### Effect of increased FR on constitutive and herbivory-induced secondary metabolism

Effects of increased FR light ratios on secondary metabolite production, specifically volatile compounds, have been shown in plants as different as *Petunia × hybrida* (Colquhoun *et al*., 2013), *Hordeum vulgare* (Kegge et al., 2015), *Ocimum basilicum* (Carvalho et al., 2016), *Nicotiana sylvestris, Solanum lycopersicon*, (Izaguirre *et al*., 2006; Cortés *et al*., 2016) and now *S. altissima*. The previous studies suggest the involvement of a wider range of phytohormones that had been found important for the growth responses. For example, a low R: FR ratio causes downregulation in the jasmonic acid (JA) pathway in shade-intolerant plants (Leone *et al*., 2014; Fernández-Milmanda *et al*., 2020). This pathway is crucial in the induced production of defensive compounds in plants, but commonly reduces plant growth (Cipollini and Lieurance, 2012). In *Arabidopsis thaliana*, JA is repressed by low R: FR light ratios (Leone *et al*., 2014) suggesting a priority of growth over defenses. Interestingly, the increased expression of IAA signaling, induced by a low R: FR ratio, has long been known as an inhibitor of JA responses and, on a mechanistic level, may explain the differential allocation of resources into cell elongation and away from secondary metabolism (Mason and Mullet, 1990; Thornburg and Li, 1991; Mason *et al*., 1992; Dewald *et al*., 1994). Moreover, the inhibition of wound- or herbivory-induced JA signaling will also significantly impair the herbivory-mediated induction of defense-related secondary metabolites (Baldwin *et al*., 1997). This would certainly explain the aforementioned findings on the FR-induced allocation into growth away from constitutive and induced secondary metabolite production and resistance in tobacco, *Nicotiana sylvestris*, and tomato, *S. lycopersicon*, (Izaguirre *et al*., 2006; Cortés *et al*., 2016). From a functional perspective, this kind of response is likely adaptive in plant systems where competition with neighbors is substantially more impacting on plant fitness than herbivory. In systems like *S. altissima* where herbivory can be the major factor mediating competition with neighbors (Carson and Root, 2000; Uesugi and Kessler, 2013), more nuanced and integrated response to the combined perception of competitors and herbivores may suit the plants better. While this project focused on the evidence for such an integration of different types of information, it goes beyond the scope of this paper to investigate the actual resistance and plant fitness effects.

However, in the light of both, the wider functional hypothesis as well as in the light of the objective of this study, there are several remarkable induction patterns in *S. altissima*’s response to FR light and herbivory. Previous studies have been shown that FR light and damage affect the production of VOCs in plants (Colquhoun *et al*., 2013; Becker *et al*., 2015; Kegge *et al*., 2015). When *S. altissima* plants are exposed to a combination of herbivore damage and higher FR light ratios, the chemical profile becomes more like one of the plants with damage. This suggests that, different from the previous studies, in the case of *S. altissima*, secondary metabolism and its induction by herbivores are not suppressed by reduced R:FR light ratios. More importantly, the fact, that the combination of FR light supplementation (i.e. perception of a potential competitor) and herbivory induced different volatile and non-volatile secondary metabolite profiles is strong support for an information integration hypothesis. Interestingly, compounds that were mostly up-regulated by higher FR light ratios in *S. altissima* plants were the ones that are down-regulated with damage or the combined exposure to supplemented FR light and herbivory and vice versa (Fig. 5A). Non-volatile compounds have been observed to change with increased FR light exposure in other study systems (Tegelberg *et al*., 2004; Kuo *et al*., 2015). Similarly, to VOCs, the changes in non-volatiles could be related to changes in the JA pathway. In how far these differential inductions of plant secondary metabolism affect subsequent interactions with other organism such as herbivores and neighboring plans remains to be determined. Interestingly, some of the compounds up-regulated by FR light supplementation in *S. altissima* are diterpenes (Diterpene 2 and 3, Fig.5B), which are known for having functions as anti-feedants and growth inhibitors for *Solidago* herbivores (Cooper-Driver & Le Quesne 1987, Uesugi and Kessler 2016). While previous studies have found downregulation of defenses in response to the exposure to FR light (Izaguirre *et al*., 2006), this up-regulation of defense metabolites in *S. altissima* indicates a differently regulated response to competition and herbivory. This is particularly remarkable when we consider that in *S. altissima* the defensive function of induced resistance and VOC-mediated information transfer (Kalske *et al*., 2019) are only realized when plants are in close proximity so that herbivores can move freely from plant to plant and so spread the risk of damage among all members of the plant population (Rubin *et al*., 2015). In conclusion, the interaction between increased FR light exposure and herbivore damage in the volatile and non-volatile chemical profile, suggests that *S. altissima* can integrate both signals (RF and light), responds with stronger induction of defenses and clearer information encoded in HIPV emissions.

### Effect of RF on the perception of HIPVs from neighbors

Our second prediction in the information integration hypothesis went one step further and suggested that if plants can integrate the information of a perceived neighbor with the information provided by an actively feeding herbivore, plants may also be able to integrate the perceived neighbor with cues that indicate future herbivory (i.e. HIPV emitted from damaged neighboring plants). Overall *S. altissima* secondary metabolite profiles did not differ in response to the exposure to VOCs from control plants or plants exposed to FR light supplementation (Fig. 4 A and B). This is not necessarily surprising as exposure to VOCs alone has rarely been found to induce significant metabolic changes without additional damage to the leaf tissue (e.g. priming of plant responses: Kessler *et al*., 2006). For example, in *S. altissima* only about 19 compounds of the non-volatile fraction of the recorded secondary metabolites were directly inducible by VOCs from neighboring plants without additional herbivore damage (Morrell and Kessler, 2017). However, we also found several individual compounds induced by the simple exposure of the plant to neighbor HIPVs. Specifically, plants under-supplemented FR light-receiving VOCs from an undamaged control plant increase the production of four VOCs dramatically (Fig.6A). This suggests that increased FR light ratios make plants more perceptive of VOCs from neighbors. Recent studies have suggested that reduced R:FR light ratios emitted from neighboring plants are used by *Arabidopsis thaliana* as a signal for kin recognition, that mediates interactions among kin neighbors, reducing competition for resources (Crepy and Casal, 2015). The FR-induced VOC emission as well as the FR-mediated differential perception of VOCs can provide an alternative and likely more specific mechanism of kin recognition. In *S. altissima*, one individual can be surrounded by several clonal ramets, which is why the availability of information about the neighbor’s genetic relatedness would be beneficial to the receiver plant because it would allow a reduced investment into resources for competition against itself or close relatives (Semchenko *et al*., 2014).

From the nine non-volatile compounds that were affected by the treatments, the greatest increment was evident in plants that received VOCs from plants with damage (Fig.6B), indicating that *S.altissima* can detect and respond to HIPVs. However, the compounds that are up-regulated are not the same in the different exposure treatments. On one hand, plant exposure to HIPVs directly induces the production of a coumaric acid derivative, a flavonoid, and two diterpene acids; on the other hand, plants simultaneously exposed to FR and HIPVs increased the production of a chlorogenic acid derivative, one diterpene acid, a caffeic acid derivative and an unknown compound. This ultimately indicates that plants exposed to light reflected off of potentially competitive neighbors perceived and processed the information encoded in HIPVs from herbivore-attacked neighbors differently from plants that stand isolated without neighbors. More generally, our data suggest that VOCs can provide specific information about the presence as well as about the identity and herbivory status of a plant neighbor. Moreover, the data support the hypothesis that *S. altissima* plants integrate the information encoded in VOCs and herbivore damage with that of spectral information from neighbors to induce distinct changes in their metabolism. While the ecological outcomes as well as the detailed molecular mechanisms of this integration of environmental information remain to be revealed, previous studies observing herbivore-induced changes of similar magnitudes reported significant effects on population and community dynamics (Kessler and Kalske, 2018). Most importantly, the implied population and species-specific differential integration of information concluded from this study may provide an explanation for plant responses that do not have outcomes predicted by the observed phytohormonal signaling (Mertens *et al*., 2021) and can explain local adaptation (Bode and Kessler, 2012) and associational resistance dynamics (Barbosa *et al*., 2009).

## Abbreviations

FR: far red
R:FR: Red to Far Red
VOC: volatile organic compounds
JA: jasmonic acid
SA: salicylic acid
HIPV: Herbivore-induced plant volatiles

## Acknowledgements

We thank Anurag Agrawal and Robert Raguso for comments on an earlier draft of this paper and Aino Kalske for insights and preliminary experiments that generated the ideas for this project. This study was conducted at Cornell University, which is located on the tradi onal homelands of the Gayogo □ hó □ n ⍰’ (the Cayuga Nation) of the Haudenosaunee Confederacy. We acknowledge the painful history of Gayogo □ hó □ n ⍰’ dispossession and honor the ongoing connec on of Gayogo □ hó □ n ⍰’ people, past and present, to these lands and waters.

## Supplementary Data

The following supplementary data are available at JXB online.

## Author contributions

AC and AK conceived the ideas and designed the methodology; AC collected the data; AC and AK analyzed the data; AC and AK led the writing of the manuscript. Both authors contributed critically to the drafts and gave final approval for publication.

## Data availability

Raw data are available on Cornell’s eCommons (https://ecommons.cornell.edu/) data repository under the title of this publication

